# A map of transcriptional heterogeneity and regulatory variation in human microglia

**DOI:** 10.1101/2019.12.20.874099

**Authors:** Adam MH Young, Natsuhiko Kumasaka, Fiona Calvert, Timothy R. Hammond, Andrew Knights, Nikolaos Panousis, Jeremy Schwartzentruber, Jimmy Liu, Kousik Kundu, Michael Segel, Natalia Murphy, Christopher E McMurran, Harry Bulstrode, Jason Correia, Karol P Budohoski, Alexis Joannides, Mathew R Guilfoyle, Rikin Trivedi, Ramez Kirollos, Robert Morris, Matthew R Garnett, Helen Fernandes, Ivan Timofeev, Ibrahim Jalloh, Katherine Holland, Richard Mannion, Richard Mair, Colin Watts, Stephen J Price, Peter J Kirkpatrick, Thomas Santarius, Nicole Soranzo, Beth Stevens, Peter J Hutchinson, Robin JM Franklin, Daniel J Gaffney

**Affiliations:** Wellcome Trust MRC Stem Cell Institute, University of Cambridge, Cambridge, UK, CB2 0QQ; Wellcome Sanger Institute, Wellcome Genome Campus, Hinxton, Cambridgeshire, UK, CB10 1SA; Division of Neurosurgery, Department of Clinical Neurosciences, Cambridge University Hospitals, Cambridge, UK, CB2 0QQ; FM Kirby Neurobiology Center, Boston Children’s Hospital, Harvard University, Boston, USA; Howard Hughes Medical Institute, Broad Institute of Harvard and MIT, Boston, USA; EMBL-EBI, Wellcome Genome Campus, Hinxton, Cambridgeshire, CB10 1SD; Biogen, Cambridge, MA, 02142, USA; Institute of Cancer and Genomic Sciences, College of Medical and Dental Sciences, Birmingham UK, B15 2TT

## Abstract

Microglia, the tissue resident macrophages of the CNS, are implicated in a broad range of neurological pathologies, from acute brain injury to dementia. Here, we profiled gene expression variation in primary human microglia isolated from 141 patients undergoing neurosurgery. Using single cell and bulk RNA sequencing, we defined distinct cellular populations of acutely *in vivo*-activated microglia, and characterised a dramatic switch in microglial population composition in patients suffering from acute brain injury. We mapped expression quantitative trait loci (eQTLs) in human microglia and show that many disease-associated eQTLs in microglia replicate well in a human induced pluripotent stem cell (hIPSC) derived macrophage model system. Using ATAC-seq from 95 individuals in this hIPSC model we fine-map candidate causal variants at risk loci for Alzheimer’s disease, the most prevalent neurodegenerative condition in acute brain injury patients. Our study provides the first population-scale transcriptional map of a critically important cell for neurodegenerative disorders.

## Introduction

Microglia are tissue resident macrophages of the central nervous system and play critical roles in neurological immune defence, development and homeostasis (Schafer and Stevens 2015; Q. Li and Barres 2018; Salter and Stevens 2017). These highly dynamic cells are challenging to study in the laboratory and are strongly influenced by different experimental environments (Gosselin et al. 2017). Genetic studies also strongly implicate microglial dysfunction in neurodegeneration (Guerreiro et al. 2013; Jonsson et al. 2013; Tansey, Cameron, and Hill 2018; Gjoneska et al. 2015) particularly in the context of the injured brain (Johnson and Stewart 2015). Single cell transcriptomics has suggested that microglial function may vary across age, sex and brain region (Olah et al. 2018; Keren-Shaul et al. 2017; Hammond et al. 2019; Masuda et al. 2019; Mrdjen et al. 2018; Mathys et al. 2017). Previous studies have used frozen post-mortem tissue from existing brain banks or fresh surgical samples typically from restricted patient groups, typically temporal lobe resections for epilepsy or peritumoral sampling. However, variability in the post-mortem index produces substantial variation in cellular expression (Welch et al. 2019). Because of this, studies of microglial activation in humans have relied on *ex vivo* stimulation with no available data from acutely injured human brains. The challenge of sampling also means that large scale genetic studies of microglia have not been attempted to date. Population studies have demonstrated that individuals subject to mild brain trauma are 5-fold more likely to develop Alzheimer’s Disease (Mackay et al. 2019). Consequently, it is of particular importance to understand the activation of human microglia in the context of acute brain injury together with the underpinning genetic contribution to neurodegeneration.

### Characterisation microglial cell populations

Here, we describe the analysis of human microglia isolated from 141 patients undergoing a range of neurosurgical procedures **(Figure 1a)**. We recruited patients from a range of pathologies, including 51 individuals with acute brain injury (haemorrhage and trauma), who sustained substantial parenchymal injury, enabling us to observe *in vivo* microglial activation. For each individual, we isolated CD11b-positive cells and performed both single cell (SmartSeq2) (Picelli et al. 2014) and bulk RNA-seq on each individual. After QC, we retained 112 bulk RNA-seq samples, and 9,538 single cells from 129 patients (**Figure 1b**). All but three of our bulk RNA-seq samples formed a single cluster with microglia from two previous studies (Y. Zhang et al. 2016; Gosselin et al. 2017), and were distinct from both GTEx brain and BLUEPRINT monocytes (**Figure 1c**). We then compared our single cell data to public datasets of 68K PBMCs isolated from a healthy donor (Zheng et al. 2017) and 15K brain cells from 5 GTEx donors (Habib et al. 2017). A total of 8,662 cells formed a cluster with the microglia population found in GTEx samples and distinct from PBMCs **(Figure 1d)** and expressed a range of known microglial marker genes, including *P2RY12, CX3CR1* and *TMEM119*, to a high level (**Extended Data Figure 1a)**. We defined this population of cells as microglia for the remainder of our analysis. We found three less common populations of cells that closely resembled other blood cell types, including NKT cells, monocytes or B-cells that comprised 8.4%, 0.5% and 0.3% of our single cell dataset, respectively. These cell types may reflect either infiltration of immune cells as a result of blood-brain barrier breakdown or intravascular contamination within the tissue. In support of the former hypothesis, we also found that the abundance of infiltrating cells strongly correlated with patient pathology, with trauma patients in particular enriched (OR=7.6, Fisher exact test *P*=1.2×10^−155^) (**Figure 1d**). We also found a significant effect of age on the abundance of infiltrating cells (3.4% increase per year, Wald test *P*=0.014) after adjusting for all known confounding factors, which could reflect blood brain barrier degeneration over the lifespan (**Extended Data Figure 1b**).

**Figure 1.**
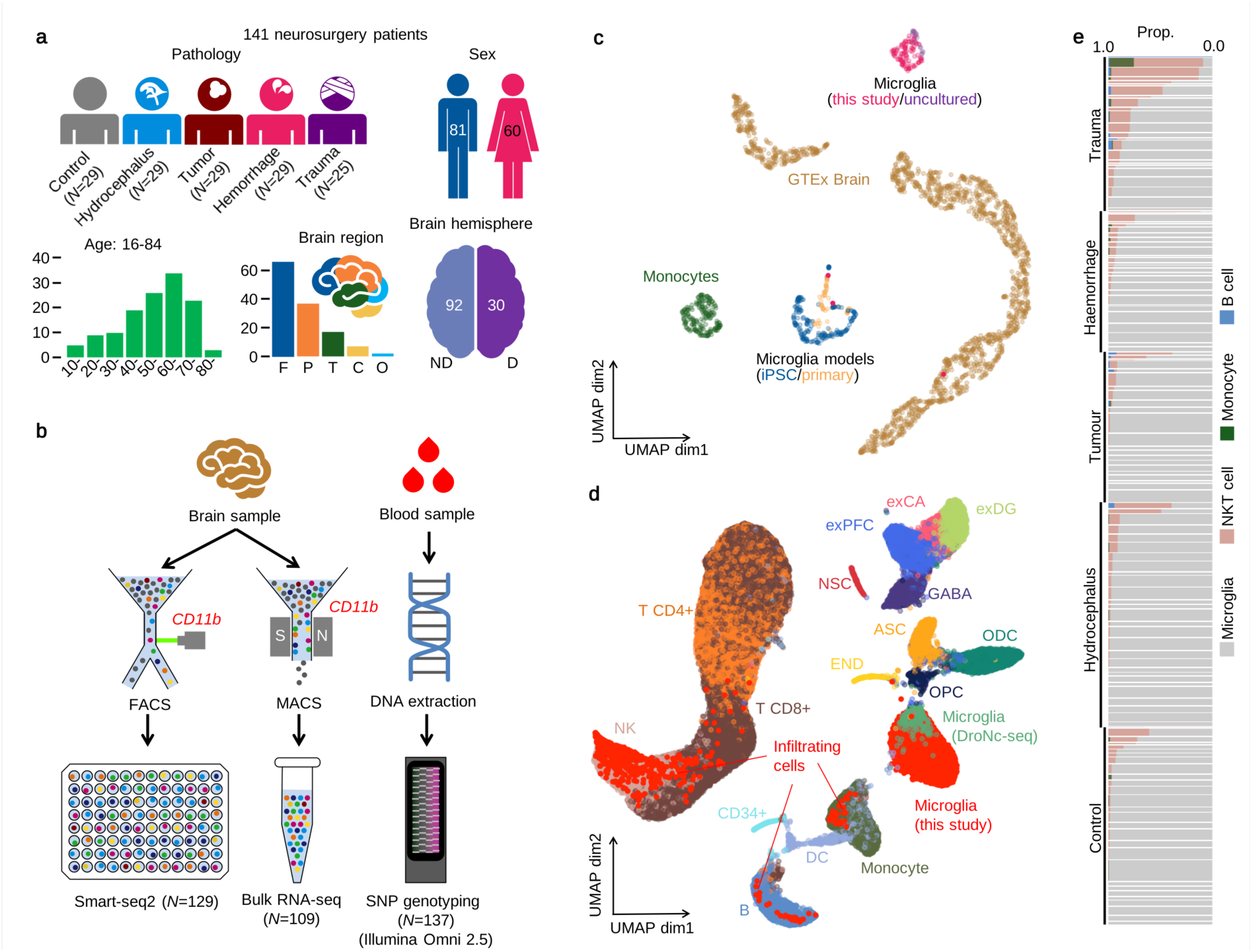
Study design and overview of the data. **a**. Metadata of 141 neurosurgery patients enrolled in this study. Brain region annotation: Cerebellum (C); Frontal (F); Occipital (O); Parietal (P); Temporal (T); non-dominant (ND); dominant (D). **b**. Experimental design using Smart-seq2 and low sample bulk RNA-seq with SNP genotyping. **c**. UMAP of bulk RNA-seq for myeloid cells and brain tissues (see Online Methods). Pink dots show our samples. **d**. UMAP of single-cell RNA-seq data combined with 68K PBMC scRNA-seq (Zheng et al. 2017) and whole brain DroNc-seq (Habib et al. 2017). Bright red dots represent cells collected in this study. Cell type annotations were obtained from: glutamatergic neurons from the PFC (exPFC); pyramidal neurons from the hip CA region (exCA); GABAergic interneurons (GABA); granule neurons from the hip dentate gyrus region (exDG); astrocytes (ASC); oligodendrocytes (ODC); oligodendrocyte precursor cells (OPC); neuronal stem cells (NSC); endothelial cells (END); dendritic cell (DC); B cell (B); hematopoietic progenitor cell (CD34+); NK T cell (NK). **e**. Proportions of non microglia for each patient in our data. Each horizontal bar corresponds to one patient. The thickness of each bar is proportional to the number of cells observed for the patient. Patients are stratified by pathology.

Within microglia, we found four subpopulations of cells (**Figure 2a)**. Two populations (C and D) were common in patients with acute brain injury (25-76% of cells) but rare in other pathologies (<5% of cells). Population B was enriched in tumor patients (OR=4.9, *P*=7.6×10^−169^) while population A was most common in control and hydrocephalus patients (**Figure 2b, c**). In populations A and B, we observed higher expression of microglial markers including P2RY12 and CX3CR1. Cells from B, C and D also demonstrated an upregulation of general immune response and cell activation (IL1B, CD83 & CCL3) (**Figure 2d, e; Extended Data Figure 2; Supplementary Table 1**). Cells from C and D exhibited additional upregulation of acute immune response pathways, including NF-kappa B, STAT3, RUNX1 as well as MHC-I expression. Population C also showed differential expression of genes associated with stress induced senescence and DNA damage (HIST1H2BG), populations D expressed genes associated with cell proliferation (FLT1) and chemotaxis (CCL4, CXCL8, CXCL16), the latter of which is shared with population B. Population B additionally showed strong upregulation of catabolic process and metabolism (GPX1) and phagocytosis (TREM2). Our cells partially overlapped with the transcriptional signatures of disease-associated microglia established in previous literature (Keren-Shaul et al. 2017; Xue et al. 2014) (**Figure 2f, g**). Taken together, these results suggest that our data contain a mix of naive microglia (population A), with three distinct states of activation that, in part, are driven by patient pathology.

**Table 1.** Colocalisation analysis of microglia eQTL with 5 AD GWAS. STH: sub-threshold; below GWAS suggestive threshold of *P*=1.0×10^−6^; NA: conditional analysis result of GWAS is not available except for Liu and Schwartzentruber (2019) (see Online Methods).

| Gene | Jansen et al. 2019 | Kunkle et al. 2019 | Lambert et al. 2013 | Marioni et al. 2018 | Liu and Schwartzentruber 2019 |
| --- | --- | --- | --- | --- | --- |
| *BIN1* | 0.984 | 0.996 | 0.994 | 0.996 | 0.996 |
| *PICALM* | 0.254 | 0.676 | 0.731 | 0.773 | 0.742 |
| *EPHA1-AS1* | 0.317 | 0.378 | 0.526 | 0.946 | 0.978 |
| *AC099524.1 (PLCG2)* | STH | STH | STH | 0.915 | 0.942 |
| *CCDC6* | STH | STH | STH | 0.685 | 0.579 |
| *ZNF646* | 0.44 | STH | STH | 0.922 | 0.664 |
| *TREML3P (tertiary)* | NA | NA | NA | NA | 0.568 |
| *SIGLEC11* | STH | STH | STH | STH | 0.99 |
| *PTK2B (secondary)* | NA | NA | NA | NA | 0.545 |
| *HLA-DRB6* | 0.062 | 0.256 | 0.517 | 0.001 | 0 |
| *HLA-DRB9* | 0.033 | 0.456 | 0.514 | 0.002 | 0 |

**Figure 2.**
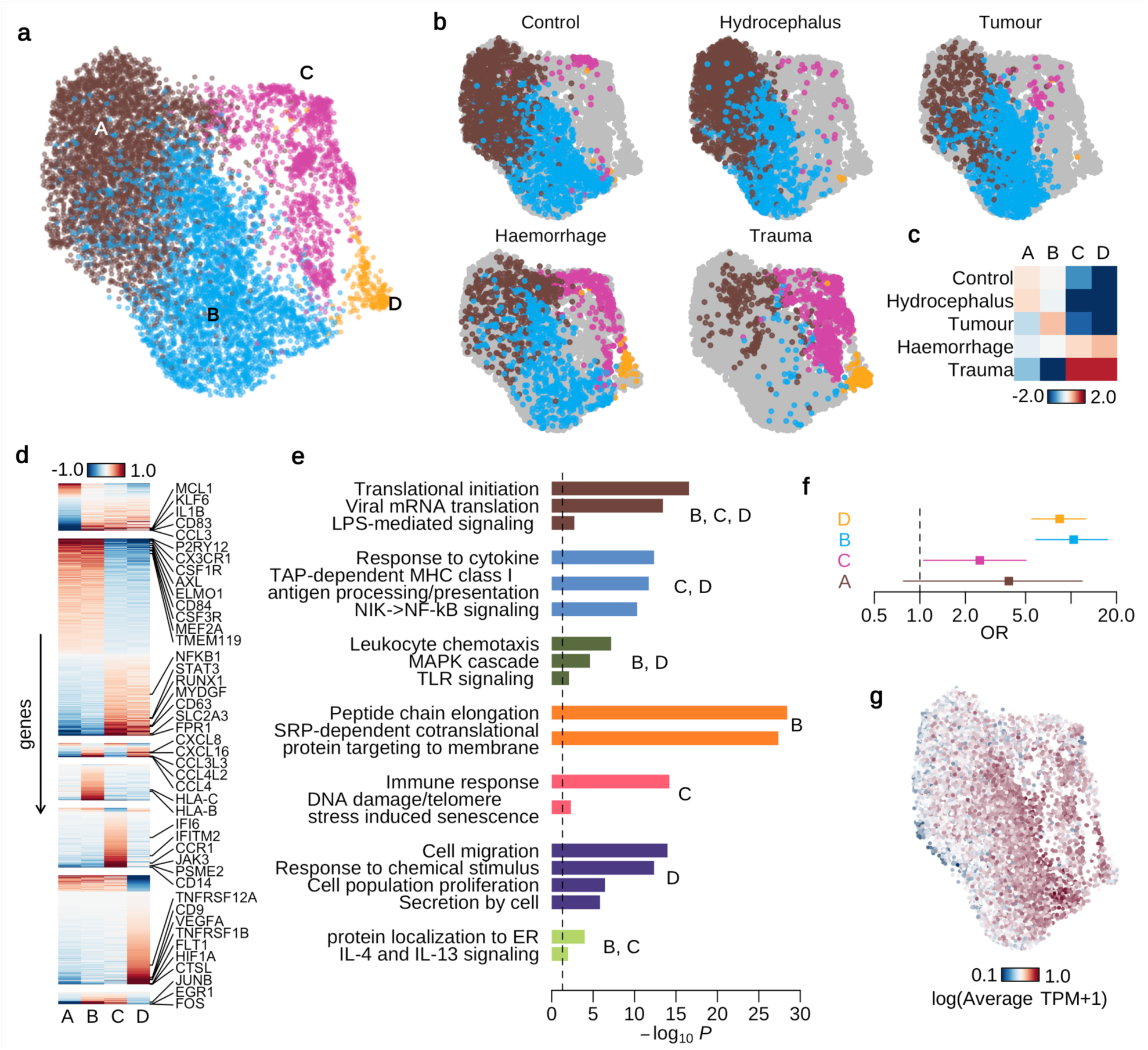
Transcriptional heterogeneity in human microglia. **a**. UMAP of 8,662 microglia cells after removing infiltrating cells. **b**. Microglial subpopulation variation between patient pathologies. Points coloured by the four different colours in Figure 2a illustrate subpopulation compositions for each pathology. Points coloured gray are all other cells. **c**. Heatmap showing the enrichment (log odds ratio) of microglial subpopulations between pathologies **d**. Heatmap of averaged, normalised expression level (defined as the posterior mean of pathology random effect term, see Online Methods) of differentially expressed genes at local true sign rate (*ltsr*) greater than 0.9 (Online Methods). Heatmap is divided into groups based on all possible pairwise groupings of the five cell subpopulations, with the most transcriptionally distinct population at the top. **e**. Pathway enrichment analysis of differentially expressed genes between different microglial subpopulations. **f**. Enrichment of disease-associated microglial (DAMs) transcriptional activation signatures in each of the four clusters in our data set. **g**. UMAP showing average normalised expression DAM genes transcriptional signatures in each cell. Red represents higher expression.

### Biological drivers of microglial expression

Our sampling design enabled us to explore the relative importance of a wide range of biological factors in driving microglial gene expression while controlling for important technical confounders, using variance components analysis. Of the biological factors we examined, clinical pathology explained more variation than all other factors combined, although all factors except sex, including age, brain region, dominant hemisphere and ethnicity, explained a fraction of variation that was significantly different from zero (LR test FWER 0.05) (**Figure 3a**). Patient explained the most variability of any single factor in the model. Although this factor captured the contribution of genetic background, it is also likely to reflect unmeasured technical effects, such as variability in cell dissociation and surgical sampling, which are confounded with patient in the model. The cellular pathways that differed between patient pathologies closely resembled the differences we observed between different subpopulations of microglia (**Extended Data Figure 3a, b**). We also detected 260 genes that varied significantly by patient age, showing upregulation of inflammation (CLEC7A, CIITA and TLR2) and downregulation of cell identity (P2RY12, CX3CR1), motility and proliferation (CSF1R) with increasing age (**Figure 3b-e; Extended Data Figure 3c; Supplementary Table 2**). Although sex explained little variation globally, we found 97 genes that were differentially expressed between males and females (**Figure 3f**). These included multiple genes in the complement pathway and synaptic pruning mechanisms (C1QA, C1QC and C3) that were more highly expressed in females than males (**Figure 3g; Extended Data Figure 3d; Supplementary Table 3**). Anatomical region of sampling also had a subtle effect on transcriptional variation, with cerebellar microglia, which are known to exhibit a distinct, less ramified morphology upregulating multiple recruitment chemokines (CCL4, CCL3, CCL4L2, CCL3L3) (**Figure 3h**; **Extended Data Figure 3e; Supplementary Table 4**).

**Figure 3.**
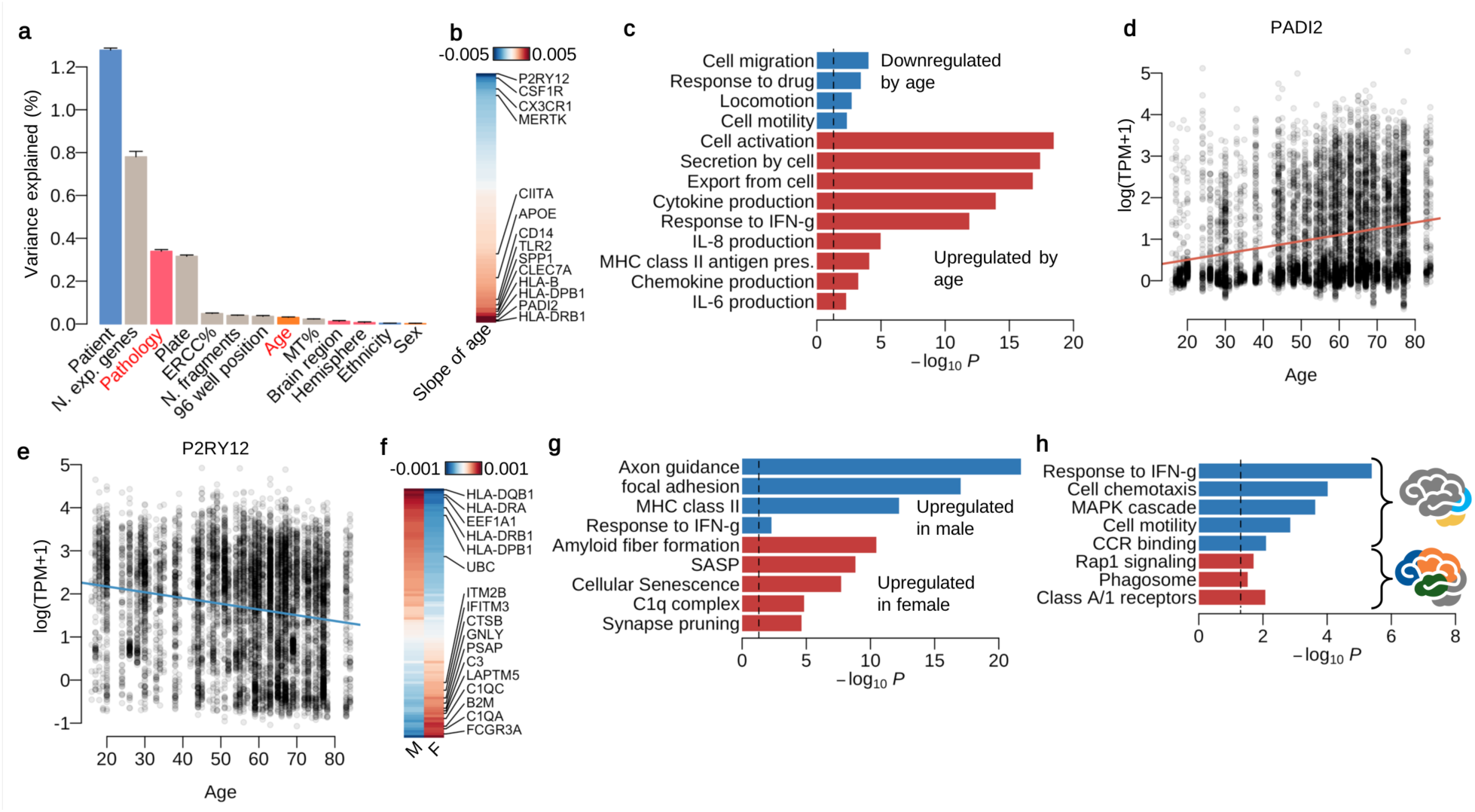
Single-cell RNA-seq revealing microglia heterogeneity driven clinical factors. **a**. Barplot shows the variance explained by each factor. Bars coloured gray are technical factors and coloured pink are clinical factors. Blue bars are partly related to patients’ genetic background. **b**. Heatmap showing the strength and direction of the age effect for differentially expressed (DE) genes at local true sign rate (*ltsr*) greater than 0.9 (Online Methods). **c**. Pathway enrichment for DE genes by age. Pathways coloured red are enriched only for DE genes upregulated by age, and the pathways coloured blue are enriched for DE genes downregulated by age. **d-e**. Example genes upregulated or downregulated by age. **f**. Heatmap showing the average normalised expression levels for DE genes by sex at *ltsr* greater than 0.9 (Online Methods). **g**. Pathway enrichment by sex. **h**. Pathway enrichment for combinations of brain regions. The blue bars show pathways upregulated in cerebellum and occipital lobe. The red bars show pathways upregulated in frontal, parietal and temporal lobes.

### eQTL mapping in human microglia and neurodegeneration

We constructed a map of expression quantitative trait loci (eQTLs) in primary human microglia. After excluding samples with low genotyping quality or substantial non-European ancestry, we mapped eQTLs using our bulk RNA-seq data from 93 individuals, and detected 401 eQTLs, summing over hierarchical model posteriors (585 eQTLs at FDR 5% using linear model). The low number of eQTLs reflected the high between-sample heterogeneity in microglia, compared with other cell types (**Extended Data Figure 4a**). We tested for colocalization of risk loci from 18 genome wide association studies (GWAS) with microglia eQTLs (**Figure 4a**), including five previous studies of Alzheimer’s disease (AD), and our own meta-analysis of these five studies for comparison (Online Methods). Across all AD GWAS, we found up to 11 risk loci with a posterior probability of colocalisation (PP4) greater than 0.5 (**Table 1**). These included well-known AD loci, such as BIN1, and less well-studied AD associations, for example EPHA1-AS1. We repeated the analysis using microglia eQTLs mapped by RASQUAL to support the colocalisation result using the allele specific expression signature (**Supplementary Table 5**). This analysis retrieved colocalisations at other AD GWAS loci, such as CD33 and CASS4. However, the test statistics may be inflated due to the additional overdispersion in our data set (**Extended Data Figure 4a**).

**Figure 4.**
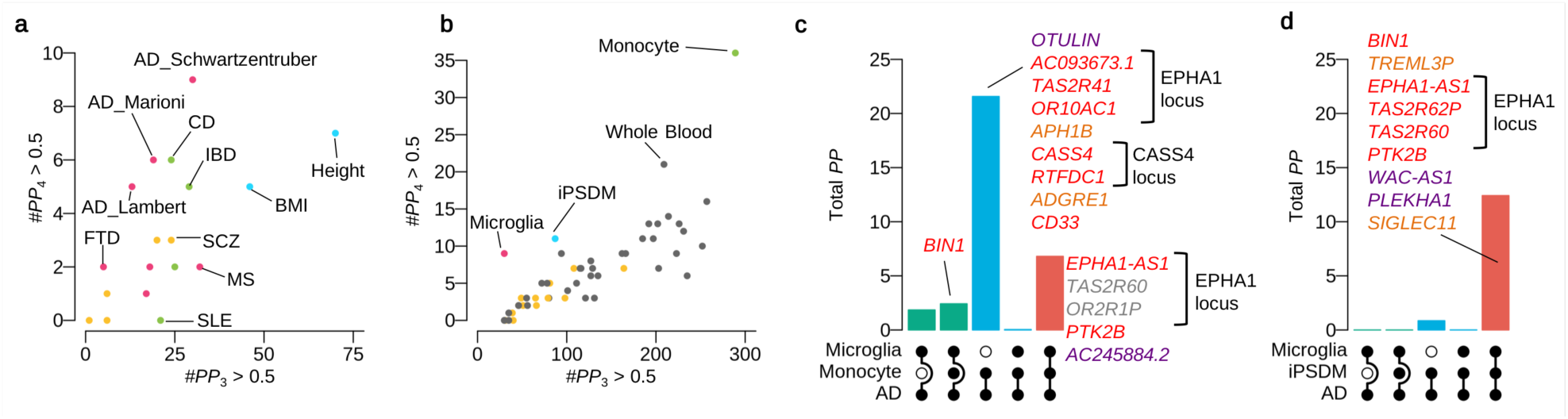
Shared genetic architecture between microglia, other myeloid cell types and human complex traits. **a**. Colocalisation of microglia eQTLs with various GWAS traits. The x-axis shows the sum of posterior probability (PP) for independent causal variants for the microglia eQTL and a GWAS trait. The y-axis is the sum of posterior probabilities for colocalised eQTLs with the trait. **b**. Colocalisation of AD GWAS (Liu and Schwartzentruber 2019) with various cell and tissues. GTEx brain tissues are coloured yellow. **c**. The number of shared eQTL genes between microglia and monocyte with AD GWAS. Genes with posterior probability greater than 0.5 for each category are listed. Each gene is coloured by the statistical significance of the lead AD GWAS variant within the 1M cis window: P<5.0×10-8 (red); 5.0×10-8×003C;=P×003C;1.0×10-6 (orange); P×003E;=1.0×10-6 (purple). **d**. Numbers of shared eQTL genes between microglia and iPSC-derived macrophage (IPSDM) with AD GWAS.

Next, we compared AD risk loci from our meta-analysis with eQTLs from the GTEx project (v7), in circulating blood monocytes and in a novel dataset of IPSC-derived macrophages (IPSDMs) from 133 healthy individuals (Online Methods) (**Figure 4b**). We found more colocalised AD eQTLs in microglia than in any GTEx brain region. We also observed many AD risk loci that colocalised with eQTLs in blood monocytes and IPSDMs. To explore the level of cell-type specificity, we mapped eQTLs jointly analysing data from microglia, monocytes and IPSDMs using a three-way Bayesian hierarchical model (**Extended Data Figure 4a, b**; Online Methods). Using this approach, we discovered 855 eQTLs, of which 108 were microglia-specific, 449 were found in all three cell types, and 192 were shared with IPSDMs but not monocytes. We also used the three-way model to evaluate the extent of sharing between the microglia eQTLs, IPSDM or monocyte eQTLs, and AD risk loci. Many colocalised AD loci, including BIN1, are found in both microglia and IPSDMs, but absent in monocytes (**Figure 4c, d**). There were also multiple AD loci where an eQTL was only detectable in circulating monocytes (e.g., CASS4 locus), although this is likely to reflect primarily the differences in power between the monocyte (n=193) and microglia data sets.

IPS models of AD risk loci are an invaluable resource for the development of future therapeutics. We next identified three AD association signals (BIN1, the EPHA1 locus and PTK2B) that colocalised with both microglia and IPSDM eQTLs (**Figure 4d**). The association for EPHA1-AS1 was shared across many cell types (**Extended Data Figure 5a, b)**, while the direction of effect at the PTK2B locus was inconsistent (**Extended Data Figure 5c)**. To fine-map causal variants we generated ATAC-seq data from 5 primary microglia and 89 IPSDMs. Colocalisation analysis revealed that the AD association signal at BIN1 was highly cell type specific (**Figure 5a**). The lead SNP of this association signal, rs6733839:C>T, was located in a region of open chromatin in both microglia and IPSDMs in which the AD risk allele. rs6733839:C>T was also associated with a significant change in chromatin accessibility (**Figure 5b**, *P*<6.1×10^−10^), and the association signal for chromatin also colocalised (PP4=0.996) with the AD association signal (**Figure 5c-f**). The AD risk allele at rs6733839:C>T created a predicted high-affinity binding site for the MEF2A transcription factor (**Extended Data Figure 5d**). We found that, although BIN1 and MEF2A are broadly expressed in many tissues, co-expression of both genes was found only in primary microglia and IPSDMs (**Extended Data Figure 5e**).

**Figure 5.**
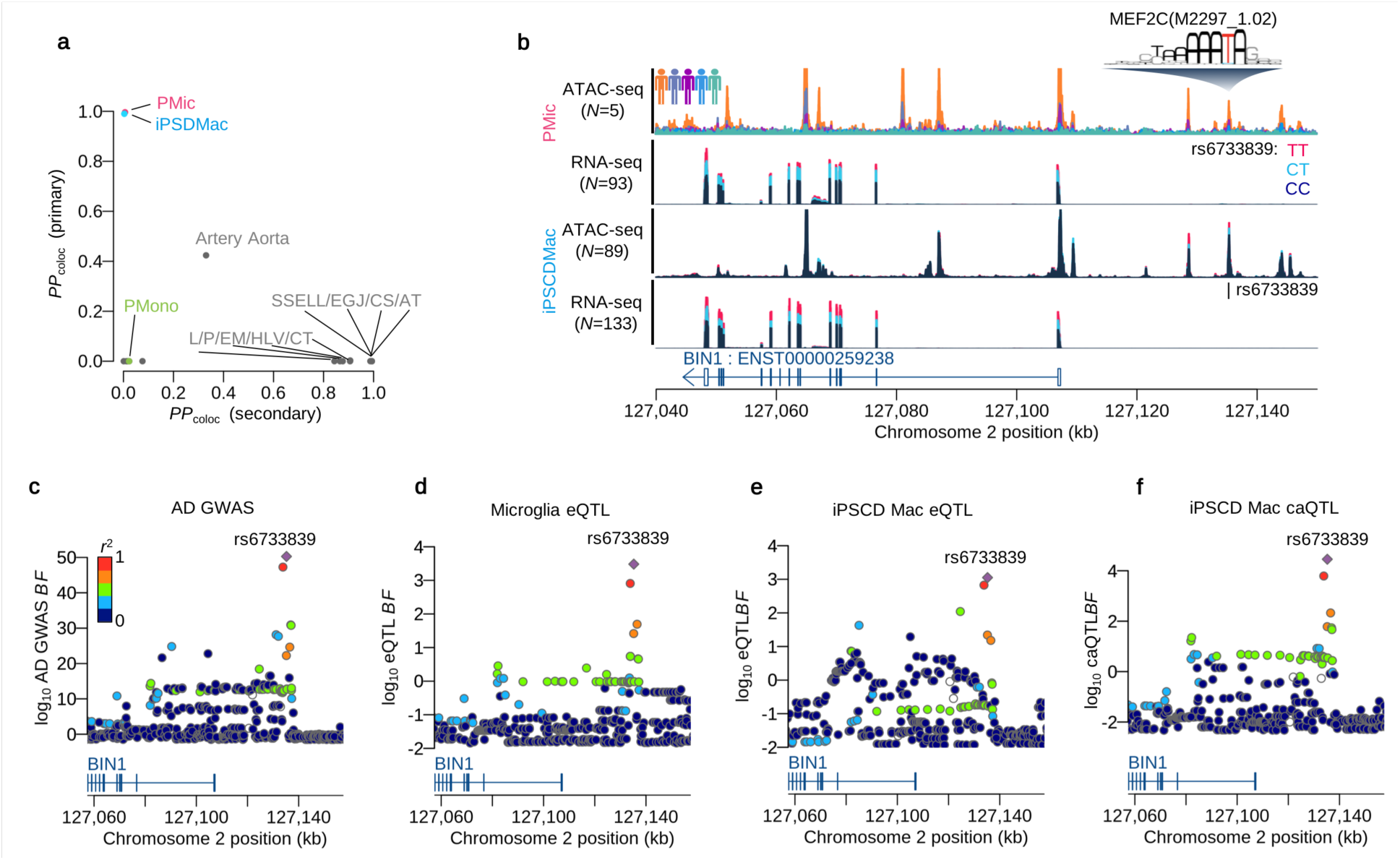
Fine-mapping of BIN1 locus. **a**. Posterior probability of colocalisation with Alzheimer’s disease (Liu and Schwartzentruber 2019). The y-axis is based on the GWAS primary signal of BIN1 locus and the x-axis is based on the secondary signal. **b**. Sequencing coverage depth of ATAC-seq and RNA-seq stratified by individuals or the three genotype groups at BIN1 lead eQTL SNP (rs6733839:C>T). The first two panels are obtained from the primary microglia and the bottom two panels are obtained from iPS cell derived macrophage. The MEF2A motif overlaps with the lead SNP and the alternative allele (T) increases predicted binding affinity. Tissue type annotation: Artery Tibial (AT), Esophagus Gastroesophageal Junction (EGJ), Colon Sigmoid (CS), Skin Sun Exposed Lower leg (SSELL), Heart Left Ventricle (HLV), Colon Transverse (CT), Esophagus Mucosa (EM), Pituitary (PI).

## Discussion

Here we present a population-level study of human primary microglia. By sampling cells from living donors, we defined transcriptional signatures of *in vivo* microglial activation avoiding artefacts from post mortem index and cell culture. We identified multiple microglial subpopulations and showed how these populations are shaped by insult, injury and other life history factors. We also created the first map of eQTLs in microglia, identified high confidence causal genes and variants underlying risk loci for Alzheimer’s disease, and identified a subset that replicated in a scalable IPS model system.

Our results underscore the variability between microglia cells from different individuals. One implication of the variation we observed between different patient pathologies is that the full spectrum of microglial function is not well cannot be captured by small studies of a single patient population. The most obvious example of this are the populations of activated microglia we identified that account for less than 5% of cells in non-trauma patients. Our results also provide a picture of the function of microglia following severe trauma, producing cell populations that exhibit a mixture of a proinflammatory and chemotactic phenotypes. Notably, although animal models of acute brain injury suggest rapid expansion of microglia following trauma (Vela et al. 2002), we only observed one population we identified had a proliferative phenotype, and both showed downregulation of CSF1R. Also in contrast to previous reports (Olah et al. 2018), we found relatively subtle effects of age on microglial transcription. The modest changes we did detect were consistent with increased inflammatory senescence in microglia over the lifespan. Likewise, differences in microglia expression between males and females were relatively small, although we did observe increased complement activity in females, perhaps suggesting a role for complement pathways in the higher incidence of AD in women.

Our eQTL analysis revealed a number of candidate risk genes for AD that function in microglia. This included well-known genes, such as BIN1, and a number of less well understood loci. One example we discovered was the EPHA1-AS1 locus, where AD risk appeared to be driven by a change in the expression of a long noncoding RNA, rather than the neighbouring protein-coding gene EPHA1 (**Extended Data Figure 5a, b**). We did not detect some well-known AD risk loci, such as CD33, with suspected function in myeloid cells. In the case of CD33, analysis of splicing patterns did indeed reveal a splice QTL at exon 2 (**Extended Data Figure 6a-c)**, consistent with previous studies (Raj et al. 2014). In other cases, we found strong colocalisation between AD risk loci and monocyte, but not microglia, eQTLs. While it is tempting to conclude that this reflects a monocyte-specific function, we believe it is more plausible that this reflects lower power in our microglia dataset and that, with an increased our sample size, many of these eQTLs would be found to be shared between the two cell types. One example of this is the CASS4 locus, where the minor allele frequency of the GWAS lead variant (rs6014724:A>G) was >10% in the monocyte data, but <5% in our microglia data set (**Extended Data Figure 6d**). Other examples of apparently spurious monocyte-specific signals included multiple genes, such as TAS2R41 and OR10AC1, that physically overlapped the second intron of EPHA1-AS1 (**Extended Data Figure 6e**)

Our comparative analysis identified multiple cases where colocalisation signals between AD and microglia eQTLs were robustly replicated in IPS derived macrophages. Although more complicated protocols for IPS-microglia differentiation exist, (Abud et al. 2017), our results highlight that IPS derived macrophages are a promising model system for deeper functional characterisation in specific cases. Our analysis also revealed instances where this assumption appeared not to hold. For example, although we found an eQTL for PTK2B that colocalised with AD in all three myeloid cell types, the direction of effect of the candidate causal variant differed (**Extended Data Figure 5c**).

In summary, we have generated a population-scale map of gene expression in primary human microglia, across a diverse set of clinical pathologies. We demonstrate the microglial response to an acute insult of the brain parenchyma. Our study provides a systematic exploration of microglia heterogeneity, defines a reference data set of microglial expression and provides a foundation for robust future functional studies of neurodegenerative disease mechanisms using IPSC-based models.

## Data Availability

Raw data (fastq files and CRAM files) of Smart-Seq2 and bulk RNA-seq for the primary microglia samples as well as bulk RNA-seq and ATAC-seq for the iPS cell derived macrophage are available from European phenome-Genome Archive (EGA) (Accession ID: EGAD00001005736). Summary statistics of eQTLs and caQTLs for primary microglia and iPSDM are also available from EGA (Accesssion ID: EGAD00001005736).

## Online Methods

### Tissue sampling

Human brain tissue was obtained with informed consent under protocol REC 16/LO/2168 approved by the NHS Health Research Authority. Adult brain tissue biopsies were taken from the site of neurosurgery resection for the original clinical indication. Paired venous blood was sampled at the induction of anaesthesia. Tissue was transferred to Hibernate A low fluorescence (HALF) supplemented with 1× SOS (Cell Guidance Systems), 2% Glutamax (Life Technologies), 1% P/S (Sigma), 0.1% BSA (Sigma), insulin (4g/ml, Sigma), pyruvate (220 g/ml, Gibco) and DNase 1 Type IV (40 g/ml, Sigma) on ice and transported to a dedicated CL 2 laboratory.

### Dissociation of brain tissue

Brain tissue was mechanically digested in fresh ice-cold HALF supplemented with 1× SOS (Cell Guidance Systems), 2% Glutamax (Life Technologies), 1% P/S (Sigma), 0.1% BSA (Sigma), insulin (4 g/ml, Sigma), pyruvate (220 g/ml, Gibco) and DNase 1 Type IV (40 g/ml, Sigma). The prepared mix was spun in HBSS+ (Life Technologies) at 300g for 5 mins and supernatant discarded. The digested tissue was rigorously triturated at 4°C and filtered through a 70 m nylon cell strainer (Falcon) to remove large cell debris and undigested tissue. Filtrate was spun in a 22% Percoll (Sigma) gradient with DMEM F12 (Sigma) and spun at 800g for 20 mins. Supernatant was discarded and the pellet was re-suspended in ice cold supplemented HALF.

### Fluorescence-activated cell sorting

For single cell smart sequencing, human microglia were using fluorescence-activated cell sorting. The isolated cell suspension was incubated with conjugated PE anti-human CD11b antibody (BioLegend) for 20 mins at 4°C. Cells were washed twice in ice cold supplemented HALF and stained with Helix NP viability marker. Cell sorting was performed on BD AriaIII cell sorter (Becton, Dickinson and Company, Franklin Lakes, New Jersey, US) at the University of Cambridge Cell Phenotyping Hub at Cambridge University Hospital, Cambridge, UK. Cells were either sorted into 98 well plates, prepared by the Wellcome Trust Sanger Institute for the purposes of single cell sequencing.

### Magnetic-activated cell sorting

To avoid sustained stress on microglia as a result of prolonged sorting times for bulk sequencing magnetic-activated cell sorting was performed on these cells. An isolated cell suspension of cells were incubated with anti-CD11b conjugated magnetic beads (Miltenyi) for 15 mins at 4°C. Cells were washed twice with supplemented HALF and passed through an MS column (Miltenyi). Each sample was washed three times in the column and then extracted. Samples were added to a 1.5 ml Eppendorf to which 350 µl of RNAlater (Qiagen) was added, samples were stored at −80°C prior to sequencing

### Blood preparation

DNA extraction was performed from the venous blood. 10 ml of whole blood was washed with 1% phosphate buffered saline (PBS) and layered on pancoll human (PAN biotech) and spun at 500g for 25 mins. The white cell component was extracted and transferred to a 1.5ml Eppendorf and stored as a frozen pellet at −80C prior to sequencing.

### SNP genotyping

Genomic DNA was extracted from blood using the QIAamp DNA mini and blood mini kit (Qiagen, 51104). 200 ng of gDNA was used for input for the SNP array (Infinium Omni2.5-8 v1.4 Kit) and genotyping was performed according to the manufacturer’s instructions. We discarded 3 samples that showed the genotyping call rate below 95% (see **Supplementary Table 6** for details). We downloaded VCF files from the 1000 Genomes Phase III integrated variant set from the project website (see URL) as the reference haplotype data. We then performed whole genome imputation for our genotype data by using the Beagle software (version 4.0; see URLs). We converted the genome coordinate from GRCh37 to GRCh38 using CrossMap (see URL).

### iPS cell culture and macrophage differentiation

We cultured 133 iPS cell lines from HipSci (Kilpinen et al. 2017). iPS cell culture and macrophage differentiation was carried as previously described (Alasoo et al. 2018) but with some minor modifications (see Supplementary Methods for details).

### Single cell RNA-seq of primary microglia

Single primary microglia cells were processed as previously described (Picelli et al. 2014) but with some minor modifications to the Nextera library making process: 0.5 ng of cDNA was used as input for the tagmentation process with all Nextera (Illumina, FC-121-1030) reagent volumes scaled down 100-fold. Tagmentation was quenched with 0.2 % sodium dodecyl sulphate. Libraries were amplified with KAPA HiFi (Kapa Biosystems, KK2601) with indexing primers ordered from Integrated DNA Technologies.

### Low-input bulk RNA-seq and ATAC-seq library preparation for primary microglia and iPS-derived macrophages

For RNA-seq samples, between 0.3 ng and 10 ng of bulk total RNA from primary microglia cells or iPS-derived macrophage cells was used as input for a modified Smart-seq2 library preparation (Picelli et al. 2014) (see Supplementary Methods for detailed protocol). ATAC-seq library preparation was performed as previously described (Alasoo et al. 2018). Pools of 96 libraries were sequenced over 8 lanes or 24 lanes of a HiSeq SBS v4 for RNA-seq and ATAC-seq preparations, respectively, collecting 75 bp paired-end reads.

### Bulk RNA-seq data of other myeloid cells and brain tissues

We downloaded fastq files of the bulk RNA-seq of 6 primary microglias (pMICs) and 9 iPS cell derived microglia (iMICs) (Abud et al. 2017), 10 monocyte derived macrophages (MDMs) and iPS cell derived macrophages (iMACs) (Alasoo et al. 2015), 10 iMICs, 8 MDMs and 4 pMICs (Douvaras et al. 2017), 45 pMICs (Gosselin et al. 2017), 9 iMICs and 3 pMICs (Muffat et al. 2016), 18 iMACs and 9 MDMs (H. Zhang et al. 2015), and 3 pMICs (Y. Zhang et al. 2016). See **Supplementary Table 7** for details of cell types and sample sizes. For brain tissues, we downloaded the count table of RNA-seq data for all tissues from GTEx (V7; see URL) and extracted 1,671 brain samples. We also downloaded fastq files of the BLUEPRINT monocyte RNA-seq data from EGA (see URL) and processed as same as our sample.

### Sequencing data preprocessing

All sequence data sets were aligned to human genome assembly GRCh38. We performed adapter trimming of Tn5 transposon and PCR primer sequences for our RNA-seq (both single-cell and bulk) and ATAC-seq data using skewer (Jiang et al. 2014) (version 0.1.127; see URLs) before alignment. Both Smart-seq2 and bulk RNA-seq data were aligned using STAR (Dobin et al. 2013) (version 2.5.3a; see URLs) using ENSEMBL human gene assembly 90 as the reference transcriptome. All other RNA-seq data were also aligned as same as our RNA-seq data without adapter trimming. Following alignment, we used featureCounts (Liao, Smyth, and Shi 2014) (version 1.5.3; see URLs) to count fragments for each annotated gene. The ATAC-seq data were aligned using bwa (H. Li and Durbin 2009) (version 0.7.4). We performed peak calling as described in (Kumasaka, Knights, and Gaffney 2019) by pooling all five samples.

### Smart-seq2 scRNA-seq quality controls

In total we sequenced 26,496 cells, of which 9,538 cells passed the quality control criteria: the minimum number of sequenced fragments (>10,000 autosomal fragments), the minimum number of expressed genes (>500 autosomal genes), mitochondrial fragment percentage (<20%) and the library complexity (percentage of autosomal fragment counts for the top 100 highly expressed genes<30%). We also performed demuxlet (Kang et al. 2018) to remove doublets from two different patients with different genetic background. We then performed cell type clustering with other primary single cell RNA-seq of 68k PBMCs and GTEx brain tissues (Zheng et al. 2017; Habib et al. 2017) (see below). We fit the latent factor linear mixed model in which the three different studies were treated as a random effect (see Supplementary Note Section 1 for details). We obtained the 12 latent factors which were subsequently used for UMAP clustering. We extracted 8,662 microglia cells in the UMAP plot (**Figure 1d**) for downstream analysis, which were distinct from other circulating blood cell types (such as NK T cells, Monocytes and B cells).

### Single cell RNA-seq data of PBMCs and whole brain

We downloaded the R binary object of 68k PBMCs from the github page (see URL). We processed the data using the provided R script and obtained the cell type annotation for PBMCs. We also downloaded the read count table of the brain DroNc-seq data from GTEx portal (see URL). The brain cell type annotation was obtained along with the count data. The count data from two studies were joined by gene IDs and converted into CPM (count per million) along with our primary microglia read count data.

### Variance component analysis

A linear mixed model of log(TPM+1) values across genome-wide genes (whose TPM>0 for 10% of total cells) was used to estimate the transcriptional variation. The 13 different factors (Patient, the number of expressed genes per cell, pathology, plate ID, ERCC percentage, the number of expressed genes in each cell, 96 well plate position, age of patient, mitochondria RNA percentage, brain region, brain hemisphere, ethnicity and sex) were fitted as random effects with independent variance parameters *ϕ*_*k*_^2^. The variance explained by the factor *k* was measured by the intraclass correlation *ϕ*_*k*_^2^/(1 + *ϕ*_*k*_^2^), where the other 12 factors were fixed constant. The standard error of the intraclass correlation was computed by the delta method with the standard error of the variance parameter estimator. See Supplementary Note Section 1.1 for details.

### Detection of microglia subpopulations

We used the linear mixed model to estimate the latent factors with the 13 known confounding factors (see Supplementary Note Section 1.2 for details). We used 15 first latent factors to cluster cells into subpopulations. We utilised the Shared Nearest Neighbour Clustering implemented in Seurat (version 3.0.2) with resolution parameter of 0.2 to identify the four microglia subpopulations.

### Differential expression analysis of known factors

We utilised the same linear mixed model we employed for the variance component analysis to adjust for 13 known confounding effects and the effect of four cell subpopulation (see above) as a random effect in differential expression analysis. We fit the model on gene-by-gene basis using the estimated variance parameters 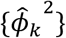 to test each factor *k* explaining a significant amount of transcription variation. If the focal factor *k* is numerical (e.g., age of patients), the Bayes factor of effect size was computed by comparing the full model and the reduced model without the factor *k*. If the focal factor *k* is a categorical variable with *l* levels (e.g., pathology with 5 levels), we partitioned the levels into any of two groups. There are 2^*l*^-1 contrasts which were tested against the null model (removing the focal factor *k* in the model) to compute Bayes factors. Then, those Bayes factors were used for fitting a finite mixture model to compute the posterior probability as well as the local true sign rate (*ltsr*) (See Supplementary Note Section 1.3 for more details). We used g:Profiler 2 implemented in R (version 2.0.1.5) to perform which pathways are enriched for DE genes for each contrast. We used genes whose *ltsr* is greater than 0.5 to perform the analysis (both upregulated and downregulated genes separately).

### Expression QTL mapping using linear regression and RASQUAL

We used simple linear regression to map eQTLs. The fragment counts were GC corrected as described in (Kumasaka, Knights, and Gaffney 2016), normalised into TPM (transcripts per million) and then log transformed (log of TPM+1). 25 principal components (PCs) were calculated and regressed out from the normalised expression levels. For each gene, We applied Benjamini-Hochberg (BH) FDR correction across all variants tested in the cis-regulatory region to obtain the minimum Q-value. Then, the minimum Q-values across all genes are adjusted again by BH FDR method to compute the genome-wide FDR. We then fit the Bayesian hierarchical model (described below) to map eQTLs using the asymptotic Bayes factors derived from the eQTL effect size and its standard error as described in (Kumasaka, Knights, and Gaffney 2019). We also mapped eQTLs using RASQUAL with the raw count data and the same 25 PCs used in linear regression as the covariates. We used --no-posterior-update option to keep the posterior genotype dosage identical to the prior genotype dosage, that allowed us to stabilise the convergence of model fitting. We picked up the minimum BH Q-value for each gene to perform the multiple testing correction genome-wide. We performed a permutation test once for each gene and constructed the empirical null distribution to which the real Q-values were compared to calibrate the FDR threshold (Kumasaka, Knights, and Gaffney 2016). Colocalisation analysis with GWAS traits was performed using the COLOC (Giambartolomei et al. 2014) implemented on R.

### Bayesian hierarchical model

We extended a standard Bayesian hierarchical model (Veyrieras et al. 2008) to jointly map eQTLs in three different cell types. We employed the association Bayes factor at each variant for each gene to compute the regional Bayes factors (RBFs) in a cis region of 1Mb centred at transcription start site (TSS) under 15 different hypotheses. Those RBFs were used in a hierarchical model to estimate prior probabilities that eQTLs are colocalised between any two of the three cell types as well as shared among three cell types. It can provide posterior probability that a gene is an eQTL for each cell type. The model also allows us to estimate the shared genetic architecture between eQTLs and GWAS loci with an appropriate prior calibration. See Supplementary Note Section 2 for more details.

### Alzheimer’s disease GWAS summary statistics

We conducted a fixed-effects meta-analysis of summary statistics from the Kunkle et al. GWAS of diagnosed AD (Kunkle et al. 2019) and a GWAS for family history of AD that we conducted in the UK Biobank (see Liu and Schwartzentruber 2019 for details) across 10,687,126 overlapping variants. For the family history of AD GWAS, we extracted AD cases from UK Biobank self-report, ICD10 diagnoses and ICD10 cause of death data, and extracted participants who reported at least one relative affected by AD/dementia as proxy-cases. We lumped the true and proxy-cases together (53,042 unique affected individuals, 355,900 controls) and performed association tests using BOLT-LMM (Loh et al., 2015). Across 36 associated loci we used GCTA to identify independently associated SNPs with a threshold of p < 10^−5^, based on LD from 10,000 randomly-sampled UK Biobank individuals. For the *BIN1* and *PTK2B* loci, we used GCTA --cojo-cond to determine summary statistics for each of the two independent signals at each locus, with a window of +/- 500 kb around each lead SNP.

## URL

GTEx: https://www.gtexportal.org/home/datasets PBMC 68k cell data: https://github.com/10XGenomics/single-cell-3prime-paper/tree/master/pbmc68k_analysis

RASQUAL (https://github.com/natsuhiko/rasqual) 1000 Genomes Phase III integrated variant set (http://www.internationalgenome.org/data) Beagle 4.0 (https://faculty.washington.edu/browning/beagle/b4_0.html) bwa 0.7.4 (https://sourceforge.net/projects/bio-bwa/files/) skewer 0.1.127 (https://github.com/relipmoc/skewer) STAR (https://github.com/alexdobin/STAR/releases) featureCounts (http://subread.sourceforge.net/)

## Extended Data Figures

**Extended Data Figure 1.**
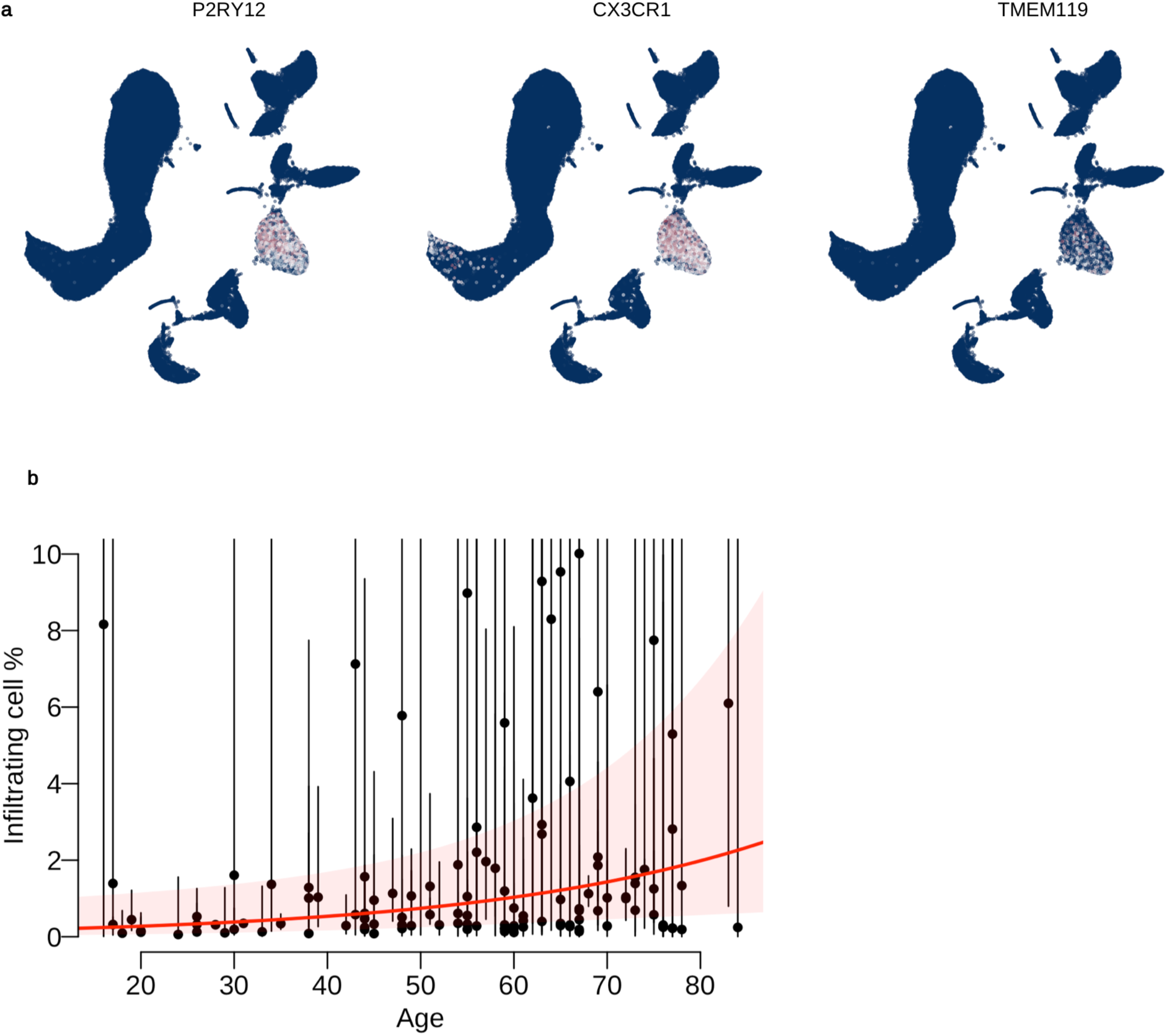
**a**. Expression of microglia marker gene, using the same UMAP coordinates as Figure 1d in the main text. **b**. Estimated age effect on the percentage of infiltrating cell percentage using the generalised linear mixed model (online Methods).

**Extended Data Figure 2.**
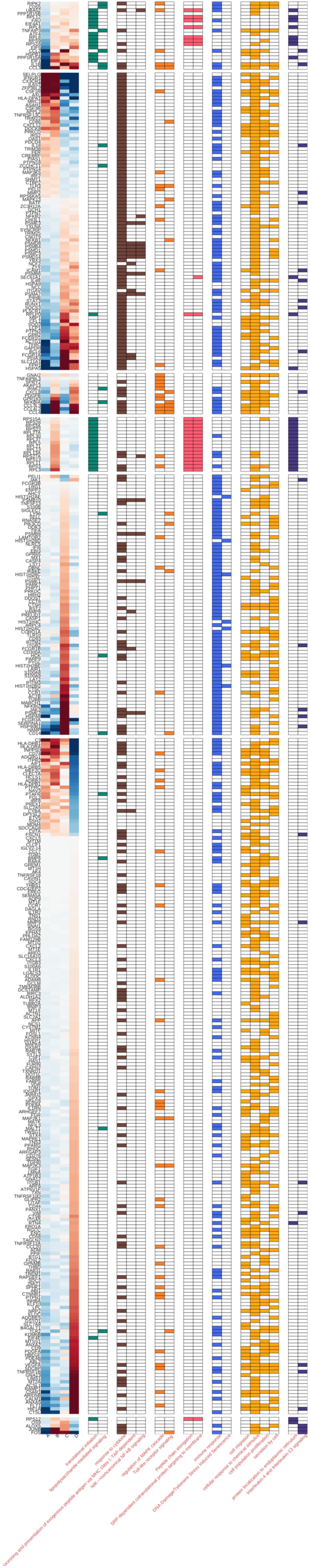
Pathways (recall>5% and precision<10%) enriched for combinations of different subpopulations.

**Extended Data Figure 3.**
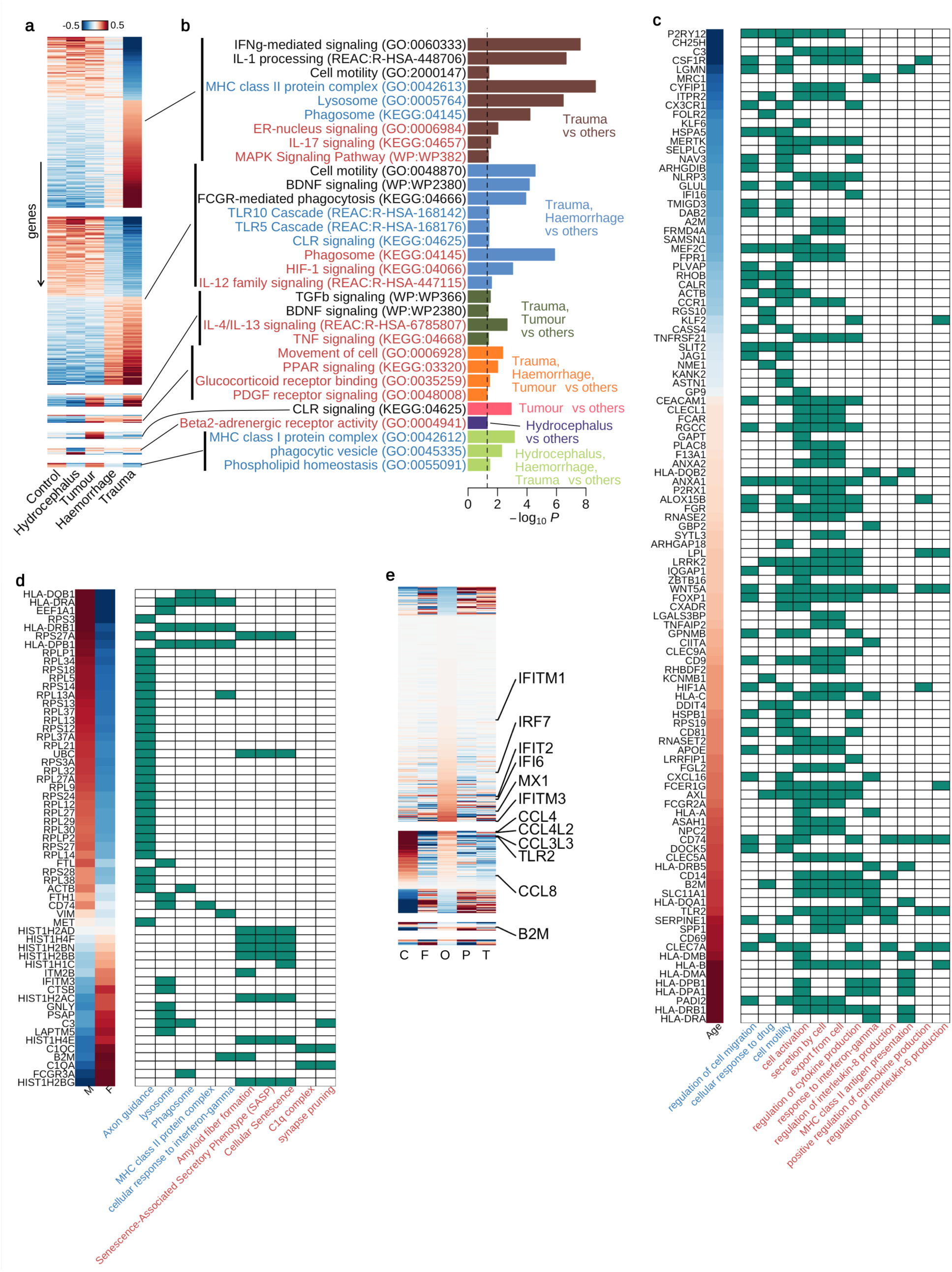
**a**. DE genes by pathology. **b**. Pathway enrichment for pathology. **c**. DE genes by age. **d**. DE genes by sex. **e**. DE genes by brain regions. C: Cerebellum; F: Frontal; O: Occipital; P: Parietal; T: Temporal.

**Extended Data Figure 4.**
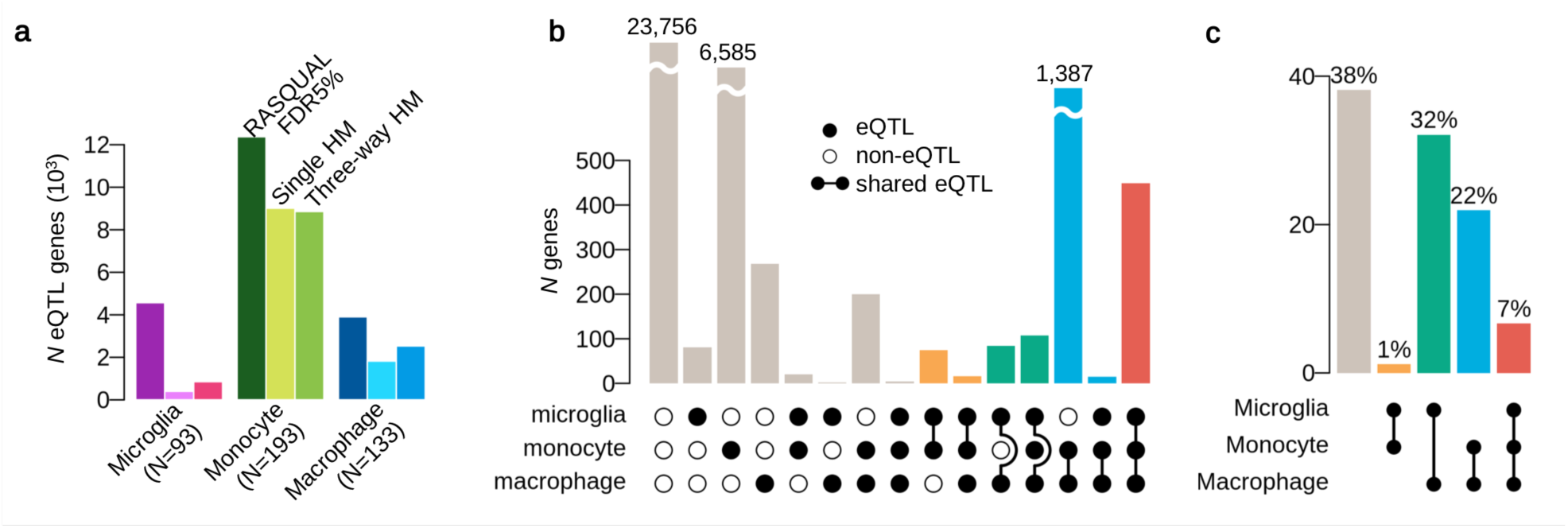
**a**. The numbers of eQTL genes discovered by three different methods for three different myeloid cell types. The left most bar is the number of eQTL genes detected by RASQUAL at FDR 5%. The middle bar is the sum of posterior probabilities derived by the simple hierarchical model (HM) and the right bar is the sum of posterior probabilities obtained by the three-way HM (see Online Methods). **b**. The number of shared eQTLs among three myeloid cell types. **c**. Empirical prior probability of eQTL sharing among three different myeloid cell types obtained by the three-way HM. The numbers are proportion of genes genome-wide and the dots connected by segment illustrates the shared genetic association.

**Extended Data Figure 5.**
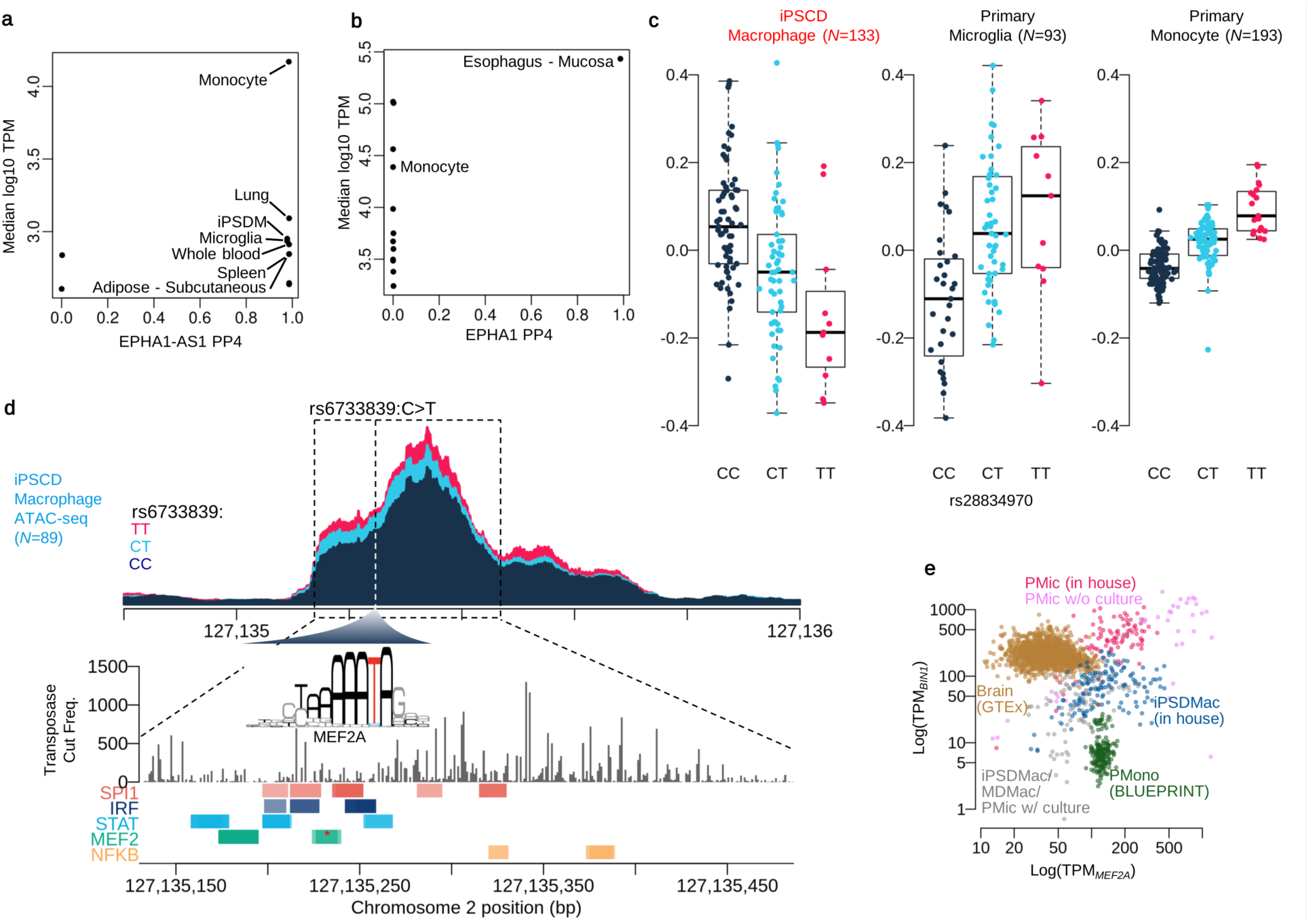
**a**. Colocalisation result between AD and EPHA1-AS1 gene for various GTEx tissues and myeloid cell types. The x-axis shows the posterior probability of colocalisation (PP4) and y-axis shows the average expression level (log10 TPM) for each tissue or cell type. **b**. Colocation result between AD and EPHA1 gene for various tissues and cells. The x-axis shows the posterior probability of colocalisation (PP4) and y-axis shows the average expression level (log10 TPM) for each tissue or cell type. **c**. Boxplots show the PTK2B eQTL effects of the lead variant (rs28834970:C>T) for the three myeloid cell types. Y-axis shows normalised expression levels (log TPM value). Each dot on the box shows the expression level of each sample. **d**. Coverage plot shows chromatin accessibility in iPS cell derived macrophages stratified by three genotype groups of the lead GWAS/eQTL variant **e**. Scatter plot of MEF2A (x-axis) and BIN1 (y-axis) expression in GTEx brain tissues and myeloid cell type.

**Extended Data Figure 6.**
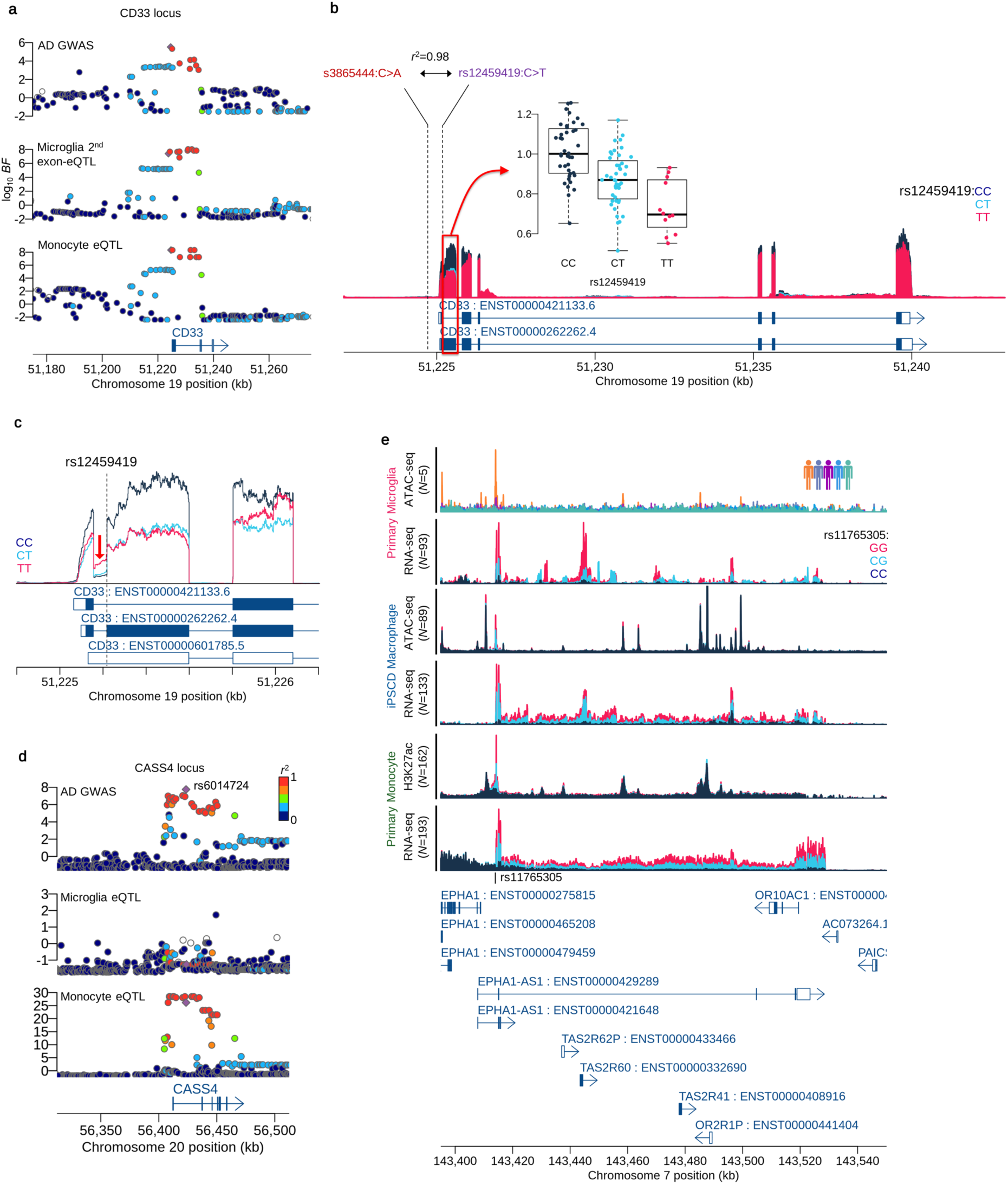
**a**. Regional plots at CD33 locus. **b**. Coverage plot shows the normalised expression level around CD33 gene stratified by three genotypes of the splicing QTL variant (rs12459419:C>T). The boxplot embedded in the panel shows splicing QTL varint effect for the 2nd exon expression (ENST0000262262.4). Each dot shows normalised expression level of each patient. **c**. Coverage plot shows expression level around the second exon (ENST0000262262.4). The coverage shows the first intron expression is negatively correlated with the second exon expression, suggesting the expression of non-coding isoform (ENST00000601785.5) is increased by the alternative allele (T) of the splicing QTL. **d**. Regional plots at CASS4 locus. **e**. Coverage plots show the chromatin accessibility, gene expression levels or histone modification ChIP-seq (H3K27ac) stratified by patients (primary microglia ATAC-seq) or three genotype groups at the EPHA1-AS1 lead variant rs11765305:G>C.

## References

Abud, Edsel M., Ricardo N. Ramirez, Eric S. Martinez, Luke M. Healy, Cecilia H. H. Nguyen, Sean A. Newman, Andriy V. Yeromin, et al. 2017. “iPSC-Derived Human Microglia-like Cells to Study Neurological Diseases.” Neuron 94 (2): 278–93.e9.

Alasoo, Kaur, Fernando O. Martinez, Christine Hale, Siamon Gordon, Fiona Powrie, Gordon Dougan, Subhankar Mukhopadhyay, and Daniel J. Gaffney. 2015. “Transcriptional Profiling of Macrophages Derived from Monocytes and iPS Cells Identifies a Conserved Response to LPS and Novel Alternative Transcription.” Scientific Reports 5 (July): 12524.

Alasoo, Kaur, Julia Rodrigues, Subhankar Mukhopadhyay, Andrew J. Knights, Alice L. Mann, Kousik Kundu, HIPSCI Consortium, Christine Hale, Gordon Dougan, and Daniel J. Gaffney. 2018. “Shared Genetic Effects on Chromatin and Gene Expression Indicate a Role for Enhancer Priming in Immune Response.” Nature Genetics 50 (3): 424–31.

Dobin, Alexander, Carrie A. Davis, Felix Schlesinger, Jorg Drenkow, Chris Zaleski, Sonali Jha, Philippe Batut, Mark Chaisson, and Thomas R. Gingeras. 2013. “STAR: Ultrafast Universal RNA-Seq Aligner.” Bioinformatics 29 (1): 15–21.

Douvaras, Panagiotis, Bruce Sun, Minghui Wang, Ilya Kruglikov, Gregory Lallos, Matthew Zimmer, Cecile Terrenoire, et al. 2017. “Directed Differentiation of Human Pluripotent Stem Cells to Microglia.” Stem Cell Reports 8 (6): 1516–24.

Giambartolomei, Claudia, Damjan Vukcevic, Eric E. Schadt, Lude Franke, Aroon D. Hingorani, Chris Wallace, and Vincent Plagnol. 2014. “Bayesian Test for Colocalisation between Pairs of Genetic Association Studies Using Summary Statistics.” PLoS Genetics 10 (5): e1004383.

Gjoneska, Elizabeta, Andreas R. Pfenning, Hansruedi Mathys, Gerald Quon, Anshul Kundaje, Li-Huei Tsai, and Manolis Kellis. 2015. “Conserved Epigenomic Signals in Mice and Humans Reveal Immune Basis of Alzheimer’s Disease.” Nature 518 (7539): 365–69.

Gosselin, David, Dylan Skola, Nicole G. Coufal, Inge R. Holtman, Johannes C. M. Schlachetzki, Eniko Sajti, Baptiste N. Jaeger, et al. 2017. “An Environment-Dependent Transcriptional Network Specifies Human Microglia Identity.” Science 356 (6344). https://doi.org/10.1126/science.aal3222.

Guerreiro, Rita, Aleksandra Wojtas, Jose Bras, Minerva Carrasquillo, Ekaterina Rogaeva, Elisa Majounie, Carlos Cruchaga, et al. 2013. “TREM2 Variants in Alzheimer’s Disease.” The New England Journal of Medicine 368 (2): 117–27.

Habib, Naomi, Inbal Avraham-Davidi, Anindita Basu, Tyler Burks, Karthik Shekhar, Matan Hofree, Sourav R. Choudhury, et al. 2017. “Massively Parallel Single-Nucleus RNA-Seq with DroNc-Seq.” Nature Methods 14 (10): 955–58.

Hammond, Timothy R., Connor Dufort, Lasse Dissing-Olesen, Stefanie Giera, Adam Young, Alec Wysoker, Alec J. Walker, et al. 2019. “Single-Cell RNA Sequencing of Microglia throughout the Mouse Lifespan and in the Injured Brain Reveals Complex Cell-State Changes.” Immunity 50 (1): 253–71.e6.

Jansen, Iris E., Jeanne E. Savage, Kyoko Watanabe, Julien Bryois, Dylan M. Williams, Stacy Steinberg, Julia Sealock, et al. 2019. “Genome-Wide Meta-Analysis Identifies New Loci and Functional Pathways Influencing Alzheimer’s Disease Risk.” Nature Genetics 51 (3): 404–13.

Jiang, Hongshan, Rong Lei, Shou-Wei Ding, and Shuifang Zhu. 2014. “Skewer: A Fast and Accurate Adapter Trimmer for next-Generation Sequencing Paired-End Reads.” BMC Bioinformatics 15 (June): 182.

Johnson, Victoria E., and William Stewart. 2015. “Traumatic Brain Injury: Age at Injury Influences Dementia Risk after TBI.” Nature Reviews. Neurology

Jonsson, Thorlakur, Hreinn Stefansson, Stacy Steinberg, Ingileif Jonsdottir, Palmi V. Jonsson, Jon Snaedal, Sigurbjorn Bjornsson, et al. 2013. “Variant of TREM2 Associated with the Risk of Alzheimer’s Disease.” The New England Journal of Medicine 368 (2): 107–16.

Kang, Hyun Min, Meena Subramaniam, Sasha Targ, Michelle Nguyen, Lenka Maliskova, Elizabeth McCarthy, Eunice Wan, et al. 2018. “Multiplexed Droplet Single-Cell RNA-Sequencing Using Natural Genetic Variation.” Nature Biotechnology 36 (1): 89–94.

Keren-Shaul, Hadas, Amit Spinrad, Assaf Weiner, Orit Matcovitch-Natan, Raz Dvir-Szternfeld, Tyler K. Ulland, Eyal David, et al. 2017. “A Unique Microglia Type Associated with Restricting Development of Alzheimer’s Disease.” Cell 169 (7): 1276–90.e17.

Kilpinen, H., A. Goncalves, A. Leha, V. Afzal, K. Alasoo, S. Ashford, S. Bala, et al. 2017. “Common Genetic Variation Drives Molecular Heterogeneity in Human iPSCs.” Nature 546 (7658): 370–75.

Kumasaka, Natsuhiko, Andrew J. Knights, and Daniel J. Gaffney. 2016. “Fine-Mapping Cellular QTLs with RASQUAL and ATAC-Seq.” Nature Genetics 48 (2): 206–13.

Kumasaka, Natsuhiko. 2019. “High-Resolution Genetic Mapping of Putative Causal Interactions between Regions of Open Chromatin.” Nature Genetics 51 (1): 128–37.

Kunkle, Brian W., Benjamin Grenier-Boley, Rebecca Sims, Joshua C. Bis, Vincent Damotte, Adam C. Naj, Anne Boland, et al. 2019. “Genetic Meta-Analysis of Diagnosed Alzheimer’s Disease Identifies New Risk Loci and Implicates Aβ, Tau, Immunity and Lipid Processing.” Nature Genetics 51 (3): 414–30.

Lambert, J. C., C. A. Ibrahim-Verbaas, D. Harold, A. C. Naj, R. Sims, C. Bellenguez, A. L. DeStafano, et al. 2013. “Meta-Analysis of 74,046 Individuals Identifies 11 New Susceptibility Loci for Alzheimer’s Disease.” Nature Genetics 45 (12): 1452–58.

Liao, Yang, Gordon K. Smyth, and Wei Shi. 2014. “featureCounts: An Efficient General Purpose Program for Assigning Sequence Reads to Genomic Features.” Bioinformatics 30 (7): 923–30.

Li, Heng, and Richard Durbin. 2009. “Fast and Accurate Short Read Alignment with Burrows-Wheeler Transform.” Bioinformatics 25 (14): 1754–60.

Li, Qingyun, and Ben A. Barres. 2018. “Microglia and Macrophages in Brain Homeostasis and Disease.” Nature Reviews. Immunology 18 (4): 225–42.

Liu, Jimmy Z., and Jeremy Schwartzentruber. 2019. Alzheimer’s Disease Meta-Analysis of Kunkle et Al GWAS and UK Biobank GWAX https://doi.org/10.5281/zenodo.3531493.

Mackay, Daniel F., Emma R. Russell, Katy Stewart, John A. MacLean, Jill P. Pell, and William Stewart. 2019. “Neurodegenerative Disease Mortality among Former Professional Soccer Players.” The New England Journal of Medicine, October. https://doi.org/10.1056/NEJMoa1908483.

Marioni, Riccardo E., Sarah E. Harris, Qian Zhang, Allan F. McRae, Saskia P. Hagenaars, W. David Hill, Gail Davies, et al. 2018. “GWAS on Family History of Alzheimer’s Disease.” Translational Psychiatry 8 (1): 99.

Masuda, Takahiro, Roman Sankowski, Ori Staszewski, Chotima Böttcher, Lukas Amann, Sagar, Christian Scheiwe, et al. 2019. “Spatial and Temporal Heterogeneity of Mouse and Human Microglia at Single-Cell Resolution.” Nature 566 (7744): 388–92.

Mathys, Hansruedi, Chinnakkaruppan Adaikkan, Fan Gao, Jennie Z. Young, Elodie Manet, Martin Hemberg, Philip L. De Jager, Richard M. Ransohoff, Aviv Regev, and Li-Huei Tsai. 2017. “Temporal Tracking of Microglia Activation in Neurodegeneration at Single-Cell Resolution.” Cell Reports 21 (2): 366–80.

Mrdjen, Dunja, Anto Pavlovic, Felix J. Hartmann, Bettina Schreiner, Sebastian G. Utz, Brian P. Leung, Iva Lelios, et al. 2018. “High-Dimensional Single-Cell Mapping of Central Nervous System Immune Cells Reveals Distinct Myeloid Subsets in Health, Aging, and Disease.” Immunity 48 (2): 380–95.e6.

Muffat, Julien, Yun Li, Bingbing Yuan, Maisam Mitalipova, Attya Omer, Sean Corcoran, Grisilda Bakiasi, et al. 2016. “Efficient Derivation of Microglia-like Cells from Human Pluripotent Stem Cells.” Nature Medicine 22 (11): 1358–67.

Olah, Marta, Ellis Patrick, Alexandra-Chloe Villani, Jishu Xu, Charles C. White, Katie J. Ryan, Paul Piehowski, et al. 2018. “A Transcriptomic Atlas of Aged Human Microglia.” Nature Communications 9 (1): 539.

Picelli, Simone, Omid R. Faridani, Asa K. Björklund, Gösta Winberg, Sven Sagasser, and Rickard Sandberg. 2014. “Full-Length RNA-Seq from Single Cells Using Smart-seq2.” Nature Protocols 9 (1): 171–81.

Raj, Towfique, Katie J. Ryan, Joseph M. Replogle, Lori B. Chibnik, Laura Rosenkrantz, Anna Tang, Katie Rothamel, et al. 2014. “CD33: Increased Inclusion of Exon 2 Implicates the Ig V-Set Domain in Alzheimer’s Disease Susceptibility.” Human Molecular Genetics 23 (10): 2729–36.

Salter, Michael W., and Beth Stevens. 2017. “Microglia Emerge as Central Players in Brain Disease.” Nature Medicine 23 (9): 1018–27.

Schafer, Dorothy P., and Beth Stevens. 2015. “Microglia Function in Central Nervous System Development and Plasticity.” Cold Spring Harbor Perspectives in Biology 7 (10): a020545.

Tansey, Katherine E., Darren Cameron, and Matthew J. Hill. 2018. “Genetic Risk for Alzheimer’s Disease Is Concentrated in Specific Macrophage and Microglial Transcriptional Networks.” Genome Medicine 10 (1): 14.

Vela, José Miguel, Angela Yáñez, Berta González, and Bernardo Castellano. 2002. “Time Course of Proliferation and Elimination of Microglia/macrophages in Different Neurodegenerative Conditions.” Journal of Neurotrauma 19 (11): 1503–20.

Veyrieras, Jean-Baptiste, Sridhar Kudaravalli, Su Yeon Kim, Emmanouil T. Dermitzakis, Yoav Gilad, Matthew Stephens, and Jonathan K. Pritchard. 2008. “High-Resolution Mapping of Expression-QTLs Yields Insight into Human Gene Regulation.” PLoS Genetics 4 (10): e1000214.

Welch, Joshua D., Velina Kozareva, Ashley Ferreira, Charles Vanderburg, Carly Martin, and Evan Z. Macosko. 2019. “Single-Cell Multi-Omic Integration Compares and Contrasts Features of Brain Cell Identity.” Cell 177 (7): 1873–87.e17.

Xue, Jia, Susanne V. Schmidt, Jil Sander, Astrid Draffehn, Wolfgang Krebs, Inga Quester, Dominic De Nardo, et al. 2014. “Transcriptome-Based Network Analysis Reveals a Spectrum Model of Human Macrophage Activation.” Immunity 40 (2): 274–88.

Zhang, Hanrui, Chenyi Xue, Rhia Shah, Kate Bermingham, Christine C. Hinkle, Wenjun Li, Amrith Rodrigues, et al. 2015. “Functional Analysis and Transcriptomic Profiling of iPSC-Derived Macrophages and Their Application in Modeling Mendelian Disease.” Circulation Research 117 (1): 17–28.

Zhang, Ye, Steven A. Sloan, Laura E. Clarke, Christine Caneda, Colton A. Plaza, Paul D. Blumenthal, Hannes Vogel, et al. 2016. “Purification and Characterization of Progenitor and Mature Human Astrocytes Reveals Transcriptional and Functional Differences with Mouse.” Neuron 89 (1): 37–53.

Zheng, Grace X. Y., Jessica M. Terry, Phillip Belgrader, Paul Ryvkin, Zachary W. Bent, Ryan Wilson, Solongo B. Ziraldo, et al. 2017. “Massively Parallel Digital Transcriptional Profiling of Single Cells.” Nature Communications 8 (January): 14049.

